# Signatures of innovation and selection in the extremotolerant yeast *Kluyveromyces marxianus*

**DOI:** 10.1101/2023.12.21.572915

**Authors:** Kaylee E. Christensen, Aniliese Deal, Jun-Ting Johnson Wang, Abel Duarte, Judith L. Edwards, Jasmine L.N. Goodman, Zhenzhen Ma, Sasha I. Padilla, Edyta Szewczyk, Catherine Rha, Rachel B. Brem

**Affiliations:** Department of Plant and Microbial Biology, University of California, Berkeley, Berkeley, CA, 94720; Graduate Group in Biophysics, University of California, Berkeley, Berkeley, CA, 94720; Graduate Group in Comparative Biochemistry, University of California, Berkeley, Berkeley, CA, 94720; Pacific Biosciences Research Center, University of Hawai’i at Mānoa, Honolulu, HI, 96822; Department of Biology, Stanford University, Stanford, CA, 94305

**Keywords:** Trait variation, evolution, fungi, stress tolerance, cell membrane

## Abstract

Organisms specialized to extreme environments can be the product of millions of years of evolutionary engineering and refinement. The underlying genetics can be quite distinct from the ones operating at earlier stages of trait innovation. In this work, we have developed the multi-stress resistant yeast *Kluyveromyces marxianus,* which diverged from its closest relative >20 million years ago, as a model for interspecies comparative biology and genomics. In growth assays of the *Kluyveromyces* genus, we found that *K. marxianus* exhibited unique tolerance of high heat and a subset of chemical stress conditions. We then generated and analyzed omic profiles from across the genus to find molecular features associated with– and potentially causal for – *K. marxianus* traits. Expression profiling revealed divergent lipid processing and membrane transport programs in *K. marxianus,* borne out in changes in lipid utilization in experimental assays. Sequence analyses found robust evidence for expansions in gene families in the *K. marxianus* genome, most notably among transmembrane transporters and in metabolic enzymes. In molecular-evolution tests, we identified adaptive protein variants throughout the *K. marxianus* genome among which plasma membrane transporters were over-represented. These data enable a model of the molecular mechanisms and evolutionary pressures underlying *K. marxianus* traits, including adaptive changes to transporters, lipid processing, and membrane functions mediating stress resistance.

**Significance statement:** Many traits of basic and applied interest arose long ago and manifest in the modern day as fixed in a given species; understanding how evolution built them, potentially over millions of years, remains a key challenge in the field. In this study, we report stress-resistance phenotypes that distinguish the yeast *Kluyveromyces marxianus* from its relatives, and we discover unique patterns of genetic and regulatory variation in membrane-protein genes, as well as unique properties of lipid metabolism, in this species. We propose a broadly applicable model in which evolution can tune membrane lipid composition and membrane-protein function to boost cellular fitness in challenging environments.

## Introduction

A central goal of evolutionary biology is to understand trait diversity in the natural world. In many cases, a phenotype of ecological or industrial interest defines a given eukaryotic species, distinguishing it from its relatives. Elucidating the genetic underpinnings of any such trait across species boundaries poses unique technical challenges. As a consequence, against a backdrop of landmark case studies mapping genotype to phenotype over deep trait divergences (Weiss and Brem 2019; Smith et al. 2020; Masly and Azom 2022), principles remain at a premium. What kinds of phenotypes tend to fix in species as they split from one another? What are the cellular mechanisms and the underlying genes and variants? Addressing these questions requires detailed study of ancient traits from physiology to the molecular level.

Single-celled fungi can serve as a powerful model system for evolutionary genetics (Hittinger et al. 2015). *Kluyveromyces,* a genus of budding yeasts, are well-suited to this paradigm given their ∼25-million-year divergence (Shen et al. 2018) and wide-ranging ecology. *Kluyveromyces* have been isolated from a range of terrestrial and aquatic environments (Anderson et al. 1986; Nagahama et al. 1999; Am-In et al. 2008; Araujo and Hagler 2011; Suzuki et al. 2014; Cernak et al. 2018; Williams et al. 2019; Chacón-Vargas et al. 2020; Al-Qaysi et al. 2021; Baptista et al. 2021; Avchar et al. 2022; Avchar et al. 2022; Spurley et al. 2022; Verma et al. 2023), but are best known for roles in food and beverage fermentations (Kurtzman et al. 2011), including elegant evolutionary analyses of the origin of lactases in dairy-associated lineages (Varela, Puricelli, Ortiz-Merino, et al. 2019; Friedrich et al. 2023). An extensive literature has focused on the growth of one species in the genus, *K. marxianus,* at temperatures exceeding 45°C, motivated by applications in the bioproduction industry (Lane et al. 2011; Lertwattanasakul et al. 2013; Mejía-Barajas et al. 2017; Madeira-Jr and Gombert 2018; Fu et al. 2019; Hoshida et al. 2020; Wu et al. 2020; Kosaka et al. 2022; Flores-Cosío et al. 2024). The evolutionary driver for the latter trait is unknown, though *K. marxianus* has been isolated from self-heating agricultural and domestic compost (Suzuki et al. 2014; Avchar et al. 2022; Verma et al. 2023). Likewise, cellular and genetic mechanisms underlying unique *K. marxianus* traits have remained incompletely understood (Nespolo et al. 2020); potential clues have emerged from comparison of this species to distantly related yeasts, including reports of resistance to chemical stress (Saini et al. 2017; Linder 2018; Illarionov et al. 2021) and expansions in sugar and potassium importer genes (Varela, Puricelli, Montini, et al. 2019; Papouskova et al. 2024).

We set out to use a comparative approach to trace in detail how *K. marxianus* diverged from its relatives. We first catalogued stress resistance characters that distinguished *K. marxianus* from the rest of the species of its clade. We then applied omic profiling to find unique expression programs and genomic features in *K. marxianus*. The results provided insights into the underpinnings and the evolution of the *K. marxianus* trait syndrome.

## Results

### *K. marxianus* exhibits a stress resistance syndrome unique in its genus

With the ultimate goal of understanding evolutionary innovations in *Kluyveromyces*, we first sought to survey stress resistance traits across the genus. We profiled the growth in liquid culture of four species associated with terrestrial habitats (*K. marxianus, K. lactis, K. wickerhamii*, and *K*. *dobzhanskii*) and three aquatic species (*K. aestuarii*, *K. siamensis*, and *K. nonfermentans*) in a panel of media conditions (Figure 1 and Table S1). At 42°C, *K. marxianus* grew far better than its relatives, to an extent beyond the pattern in a permissive control condition (rich medium at 28°C; Figure 1A); genetically distinct *K. marxianus* isolates from various substrates and geographical origins phenocopied each other with respect to high-temperature growth except for one thermosensitive strain isolated from a fly (Figure S2, Table S1), as expected if loss of thermotolerance were a rare and recent event in this predominantly thermotolerant species (Lane et al. 2011). In ethanol, caffeine, the DNA intercalator propidium iodide, and the DNA damage agent methyl methanesulfonate (MMS), *K. marxianus* likewise outperformed the other species of the genus (Figure 1A). Our profiling also identified other conditions in which *K. marxianus* did not stand out with respect to growth (Figure S1B). We conclude that *K. marxianus* is unique in the genus for its resistance to a broad subset of abiotic challenges, including high temperature.

**Figure 1.**
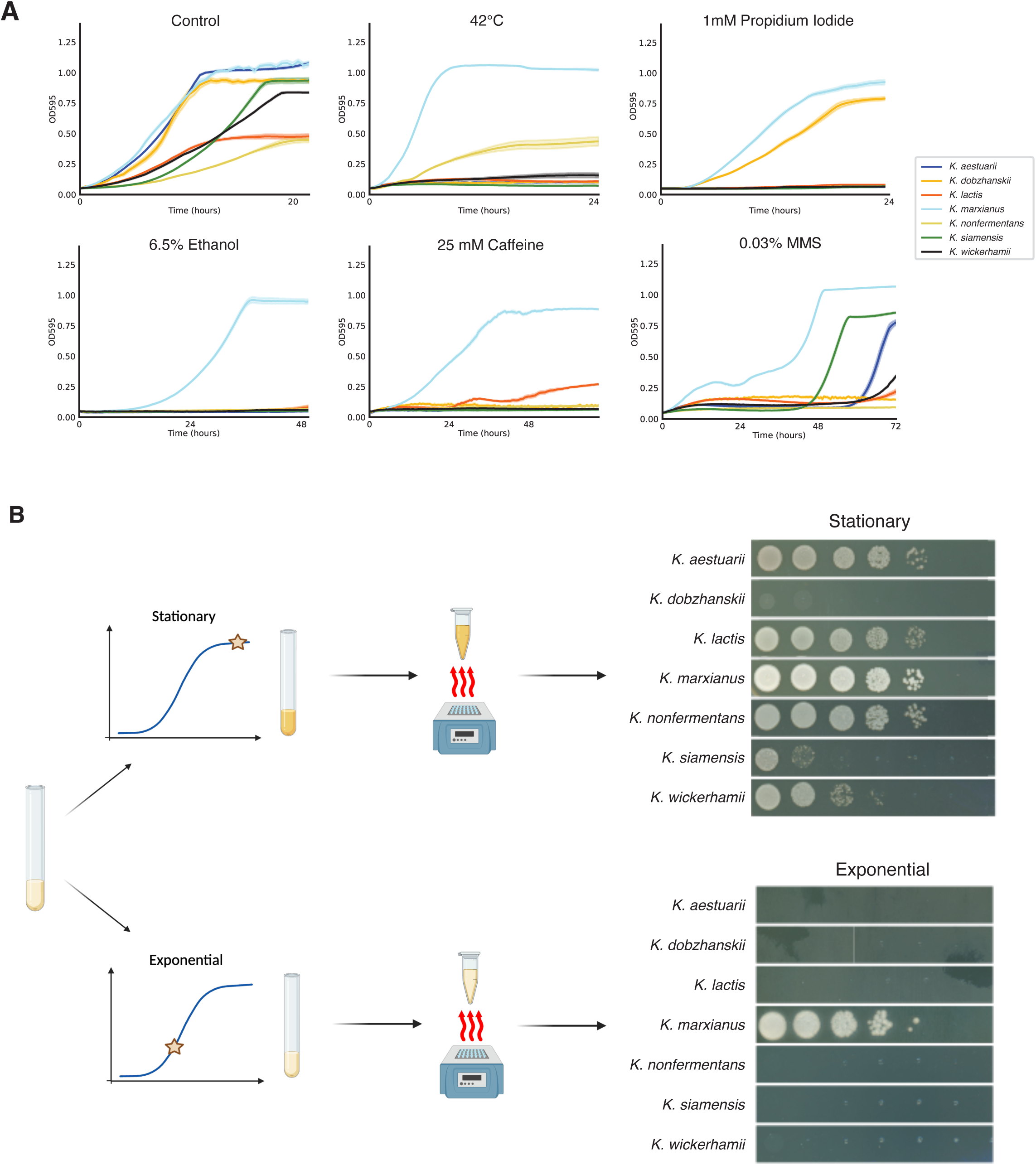
*K. marxianus* shows a unique heat and chemical stress resistance phenotype that only manifests during its growing stage. A, Each panel reports the results of a growth profiling experiment across the genus in the indicated condition. In a given panel, each trace reports a growth timecourse for the indicated species, with the solid line displaying the mean across technical replicates and the faint outline showing the standard error. OD, optical density. **B**, Each row of serial dilutions reports viability after heat shock of a sample of the indicated species in the indicated liquid growth phase. Schematics show experimental design of heat-shocking cells either at stationary or exponential phase. See Figure S3B for unshocked controls.

### The uniqueness of *K. marxianus* resistance phenotypes depends on growth phase

For many microbes, when cells face nutrient exhaustion, they shut down exponential (log-phase) growth and enter the quiescent, extra-durable state called stationary phase. The difference between *K. marxianus* and its relatives in stress resistance that we had noted across a growth timecourse (Figure 1A) might also manifest in the stationary state or, alternatively, its cellular mechanism could be tied to active cell division. To distinguish between these possibilities, we focused on thermotolerance as a test case. We used a viability assay in which we inoculated a given liquid culture into rich medium; incubated for growth at the permissive temperature until nutrient exhaustion; subjected this stationary-phase culture to a short heat shock; and used it to seed colonies for growth at the permissive temperature on solid medium (Figure 1B). Results revealed only modest differences between species in heat shock resistance under this stationary-phase heat treatment, with no particular advantage for *K. marxianus* (Figure 1B). By contrast, when we treated exponential-phase cultures with heat shock, *K. marxianus* exhibited much higher viability than did the other species of the genus (Figure 1C), paralleling its advantage during chronic high-temperature exposure (Figure 1A). No such differences were apparent in a permissive 28°C control experiment (Figure S3B). We conclude that for high temperature, the *K. marxianus* advantage only manifests when cells are actively dividing, and that cell death, rather than slow or arrested growth, mediates the defects of the rest of the genus.

To extend our analysis of cell death as the underlying cellular basis of species differences in stress, we next assayed viability in exponential-phase cultures across a panel of representative stresses, focusing on the stress-sensitive species *K. lactis* as a point of comparison for *K. marxianus* (Figure S1A). The latter exhibited a survival advantage in all stress conditions tested (Figure S4). The difference was amplified by oxygen depletion, though both species performed better in normoxia (Figure S4), as expected (Lertwattanasakul et al. 2013; Mejía-Barajas et al. 2017; Madeira-Jr and Gombert 2018; Fu et al. 2019; Hoshida et al. 2020; Kosaka et al. 2022). Together, these trends from assays of viability establish the resistance character of *K. marxianus* as a lineage-unique gain of protection against lethal failures of the proliferation program under temperature and chemical stress.

### Unique expression and lipid utilization programs in *K. marxianus*

To learn more about the underpinnings of divergence across *Kluyveromyces*, we turned to an expression profiling approach, for which we again chose *K. lactis* as a representative stress-sensitive species for comparison against *K. marxianus*. We surveyed the transcriptomes of the two species under exponential growth in representative treatments where we had observed divergent growth phenotypes — high temperature, caffeine, MMS, and ethanol — and in the permissive control of rich medium at 28°C, with stress conditions chosen to enable growth of both species (Table S2 and see Methods). Principal component analyses of normalized RNA-seq read counts made clear that the major source of variation across the data set was a marked difference between *K. lactis* and *K. marxianus* transcriptomes in all conditions, including controls (Figure 2A, Figure S5, and Table S2). Functional-enrichment tests revealed induction of membrane-lipid trafficking and intracellular transport, clathrin endocytosis, and cell wall genes driving the values of the major principal component in *K. marxianus* transcriptomes (Figure 2B and Table S2). High expression of translation and ribosomal RNA processing genes drove the values of the major principal component in *K. lactis* samples (Figure S5 and Table S2). Apart from these effects, the second principal component reflected a suite of genes that *K. lactis* turned up and down depending on treatment, whereas the *K. marxianus* transcriptome was much less volatile with respect to stress (Figure 2A, Figure S5, and Table S2), as expected if *K. lactis* underwent more functional failures during growth in our panel of conditions. Most salient in the set of stress-responsive genes in *K. lactis* was heightened induction of translation gene sets over and above the effect we had noted in control conditions (Figure 2C, Figure S5, and Table S2), dovetailing with previous studies (Lertwattanasakul et al. 2015). Functional-genomic enrichment analysis of raw expression measurements yielded similar trends (Table S3).

**Figure 2.**
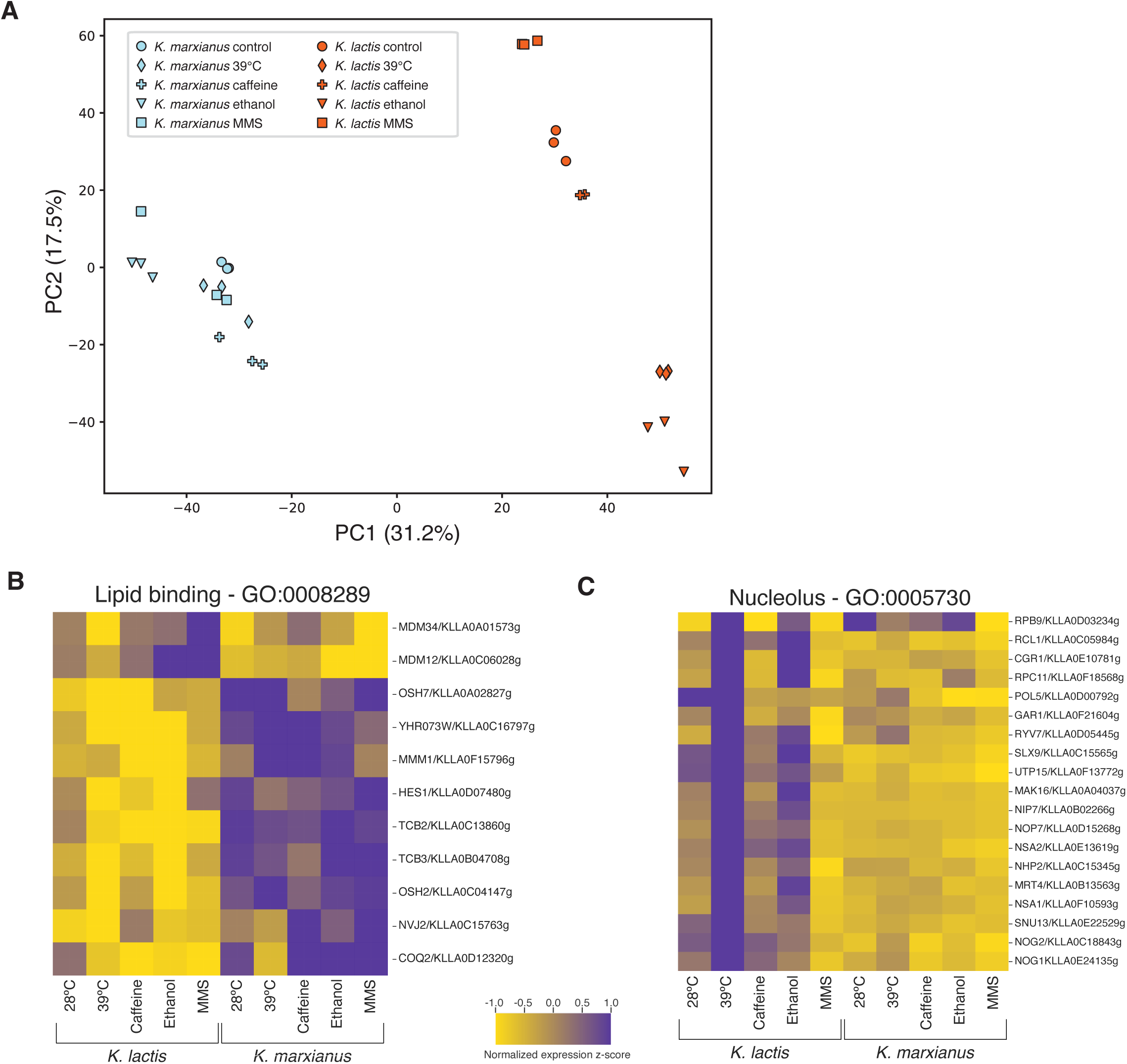
Overall regulatory volatility by *K. lactis* relative to *K. marxianus* and distinct regulatory patterns that separate the species. A, Shown are results of principal component (PC) analysis of *K. lactis* (orange) and *K. marxianus* (blue) transcriptomes across treatments. Each point reports the values of the top two PCs for one replicate transcriptome of the indicated species subjected to the indicated stress treatment or a 28°C unstressed control. In a given axis label, the value in parentheses reports the proportion of variance across the set of transcriptomes explained by the indicated PC. **B,C,** Each panel reports the effect of stress on expression of genes of the indicated Gene Ontology term in *K. marxianus* and *K. lactis*. In a given panel each cell reports, for the indicated gene, normalized read counts in the indicated stress treatment and an unstressed control, standardized by row using z-scores.

Our observations of divergent expression of membrane-lipid trafficking genes in *K. marxianus* (Figure 2B and Table S2) echoed previous comparative-biology studies of an unusual inability by *K. marxianus* to take up sterol from growth medium (Dekker et al. 2021) and unusual effects of sterol-metabolism mutations (Ren et al. 2024). We hypothesized that *K. marxianus* could likewise exhibit divergent phenotypic responses to other classes of lipid. To explore this notion, we used an experimental paradigm testing utilization by yeast of exogenous lipids when *de novo* fatty acid synthesis is blocked by the drug cerulenin (Jung et al. 2013). Using oleate as the lipid supplied in the medium in this assay, we found *K. marxianus* to be among the lowest performers of the genus (Figure S6), a surprising contrast to its advantages in stress conditions (Figure 1A). This minimal exogenous oleate processing by *K. marxianus*, even when it was required for optimal growth, paralleled the behavior of *K. marxianus* in sterols (Dekker et al. 2021), suggestive of a hard-wired preference by this species for internal lipid sources. Interestingly, the aquatic species *K. siamensis* and *K. nonfermentans* also failed to utilize exogenous oleate in our assays (Figure S6), plausibly as a product of independent evolutionary losses in their niche.

### Unique transporter and metabolic gene duplications in the *K. marxianus* genome

We reasoned that genomic features distinguishing *K. marxianus* from its relatives could serve as a source for clues to the underlying molecular basis of its traits. As a first strategy to find genomic novelties in *K. marxianus*, we focused on species-specific copy number change in paralogous gene families. From annotated reference genomes for six *Kluyveromyces* species (*K. marxianus, K. lactis, K. wickerhamii*, *K. dobzhanskii*, *K. aestuarii*, and *K. nonfermentans*; Figure S1A), plus outgroups from nearby clades in the Saccharomycotina subphylum (including *S. cerevisiae* and *C. albicans* as well-annotated model organisms), we identified rapidly changing gene families in each branch with tools of the CAFE 5 package (Mendes et al. 2021). This approach identified 16 gene families with inferred gain or loss events along the *K. marxianus* lineage at rates that were unusual relative to the average across the species set (Figure 3A and Table S4), including families subject to tandem duplications and those with genes across the *K. marxianus* chromosomes (Figure 3B and Table S4). Of these, 13 were families with expansions in *K. marxianus* that could be classified based on domain annotations as functioning in metabolism or localized to the plasma membrane, the latter including transporters annotated as nutrient importers and xenobiotic efflux pumps (Figure 3A). These data make clear that gene family expansions are prevalent in the *K. marxianus* genome, and they highlight membrane proteins and metabolism as compelling candidate molecular players underlying *K. marxianus* phenotypes.

**Figure 3.**
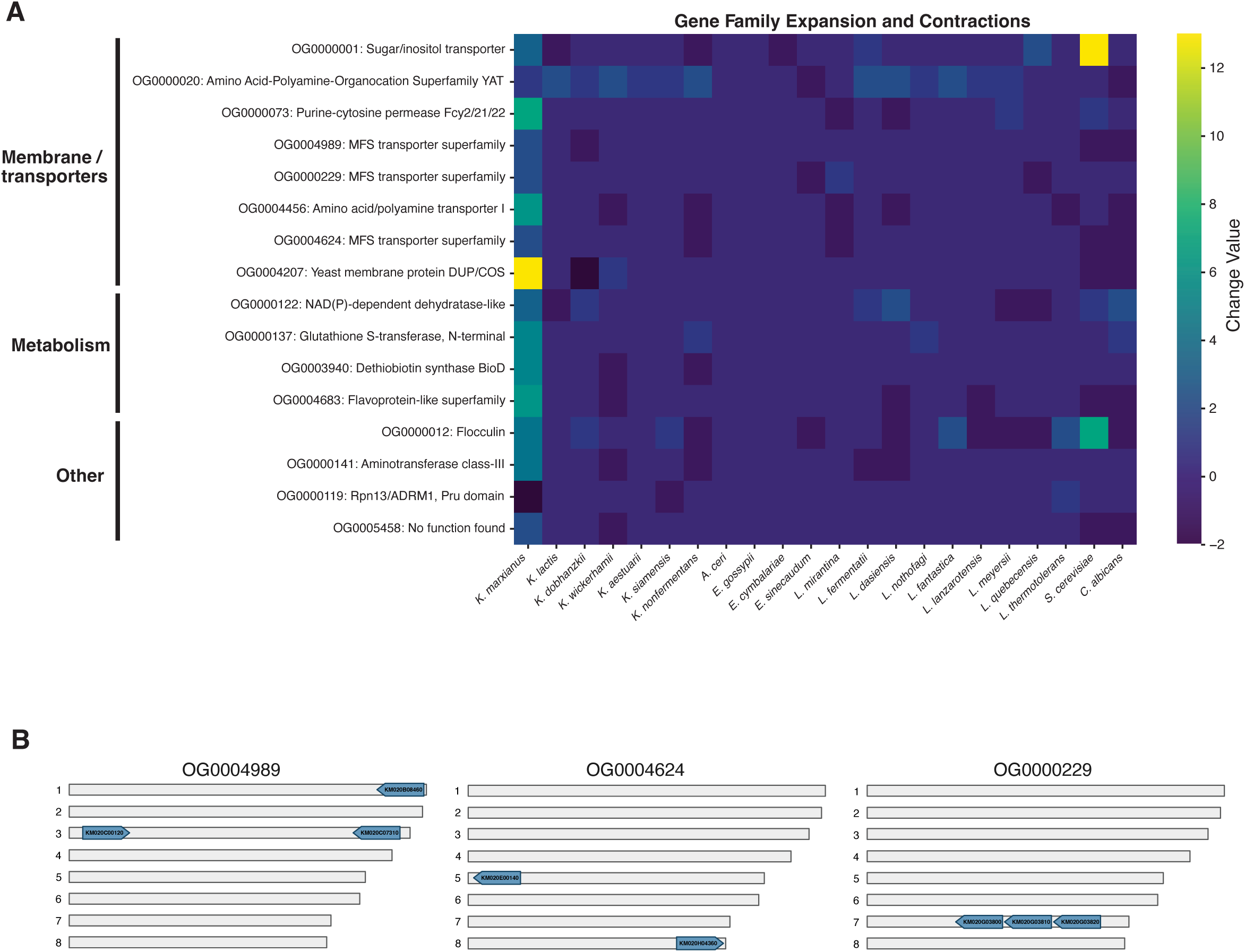
Gene family expansions in *K. marxianus*. A, The color of each cell reports the change in copy number, in the indicated genome of a *Kluyveromyces* or related species relative to its parent node in the phylogeny, of a gene family (orthogroup, OG) with significant expansion or contraction in the *K. marxianus* genome as inferred with CAFE 5. Transposon and reverse transcriptase families with significant change values are not listed, see Table S4 for results for all analyzed orthogroups. **B,** Each panel reports the positions of genes in the *K. marxianus* genome in one indicated gene family from **A** with annotations as major facilitator superfamily (MFS) transporters. In a given panel, each row represents a chromosome in *K. marxianus*, and each arrow reports a gene of the respective family and its transcriptional direction (not to scale).

### Widespread phylogenetic evidence for positive selection on amino acid alleles in *K. marxianus*

As a further means to identify genomic correlates of *K. marxianus* phenotypes, we pursued molecular-evolution analyses of amino acid variation on a per-gene basis. We reasoned that some phenotypes unique to *K. marxianus* could have evolved as a product of adaptation to challenging ecological niches, and that at relevant loci, we could detect evidence for positive selection on protein-coding sequences. For a phylogenetic test toward this end, at each gene in turn, we aligned one-to-one orthologs from reference genomes for our six focal *Kluyveromyces* species, to which we fit models in the PAML (Yang 2007) and HyPhy (Pond et al. 2005) packages earmarking codons subject to positive selection along the *K. marxianus* lineage.

Emerging from these tests were >500 and 80 genes with evidence for positive selection from PAML and HyPhy analyses, respectively (Figure 4A-B), overlapping in 51 cases (Figure 4B and Table S5). We considered the latter genes, given their signal in multiple independent analysis frameworks, as representing the strongest cases for positive selection in *K. marxianus* according to phylogenetics. This hit list included the potassium transporter *TRK1*, whose *K. marxianus* ortholog is known to be hyperactive relative to that of model yeast (Papouskova et al. 2024), and other genes annotated in transmembrane transport as well as metabolism, cell wall structure, and protein production (Figure 4B and Table S5). Control analyses revealed far less signal for positive selection on the lineage leading to *K. lactis* (Figure 4A and Table S5), supporting the inference of a history of particularly pervasive adaptation at the amino acid level in *K. marxianus*.

**Figure 4.**
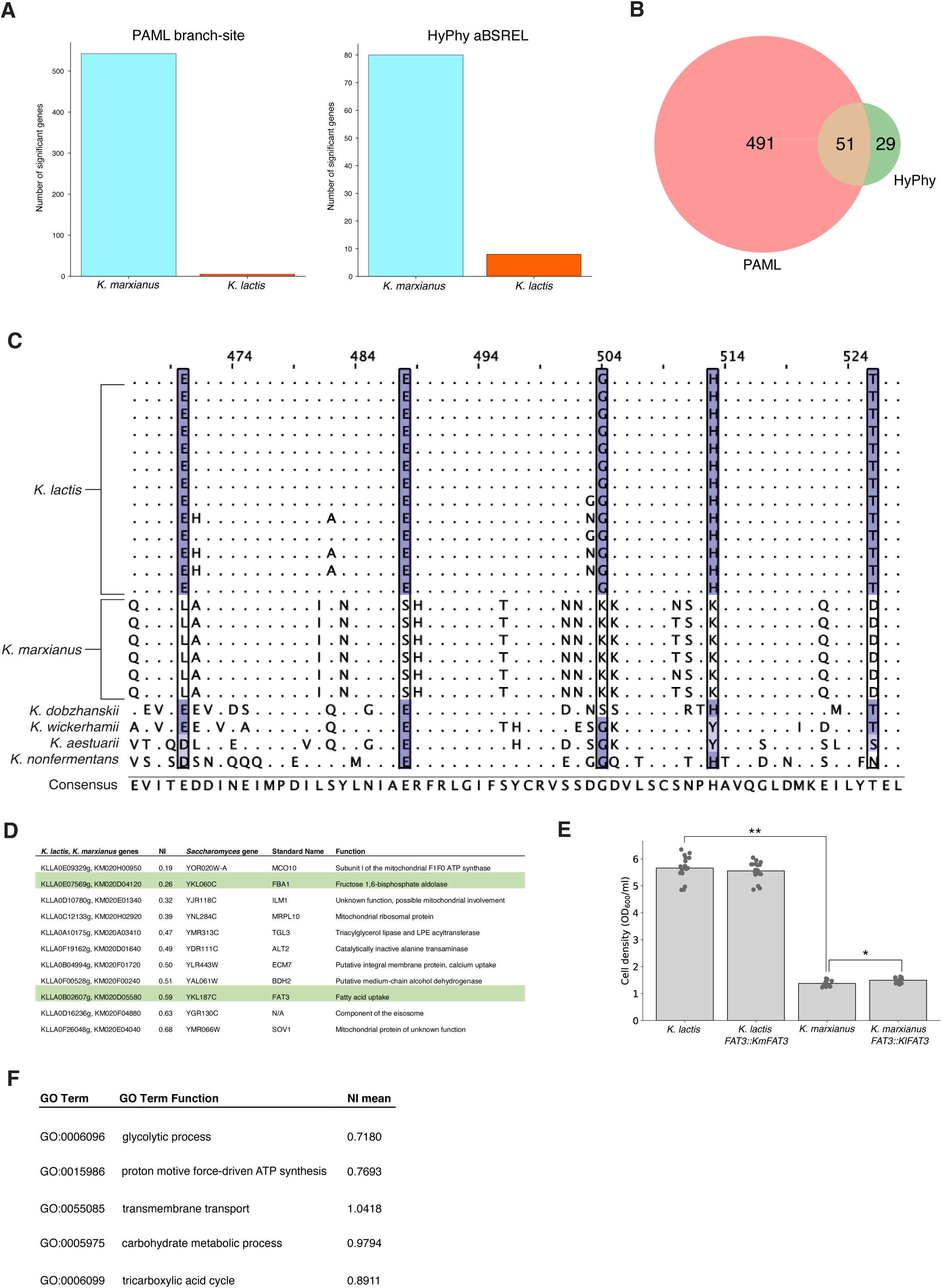
Phylogenetic analysis reveals accelerated amino acid evolution in *K. marxianus*. A, Each bar shows the number of genes with significant evidence for positive selection on the indicated lineage (adjusted *p* > 0.05) in a phylogenetic PAML branch-site test and HyPhy aBSREL test. **B,** Venn diagram of genes called significant for positive selection in *K. marxianus* in PAML branch-site and HyPhy aBSREL tests. **C,** Amino acid alignment of an example region of a gene, KLLA0B02607g (*S. cerevisiae* ortholog FAT3/*YKL187C*), with evidence for positive selection from the PAML test in **A** and from the McDonald-Kreitman test of **D**. Highlighted columns show inferred targets of positive selection from the PAML test in **B**. Highlighting indicates the amino acid’s BLOSUM 62 matrix score with respect to the consensus amino acid. **D,** Each row reports a gene with evidence for adaptive protein variation between *K. marxianus* and *K. lactis* according to the McDonald-Kreitman test (adjusted *p* < 0.05). NI, neutrality index. Genes highlighted in green also showed significant evidence for positive selection on the branch leading to *K. marxianus* in phylogenetic tests (PAML branch-site adjusted *p* < 0.05 and HyPhy aBSREL adjusted *p* < 0.05; Table S5). Not shown here are genes with *p*-value < 0.05 as a product of exceptionally high or low amino acid polymorphism; see Methods and Table S5. **E,** Each bar shows the cell density (as OD_600_/mL after 24 hours of *K. marxianus* and *K. lactis* and their respective FAT3 allele swaps in media with cerulenin and oleate, where cerulenin blocks lipid uptake and oleate is the only available lipid for utilization. The starred comparison between *K. marxianus* and its klFAT3 mutant indicate a statistically significant difference (*p* = 0.0223). **F**, Each row reports a gene ontology term with significant enrichment for low NI (adjusted *p* < 0.05) in the McDonald-Kreitman test (Table S5).

For most of the many gene hits from our phylogenetic tests focused on *K. marxianus*, a small number of codons per gene were classified as targets of positive selection (Table S5). We expected that if the derived alleles at each such site had undergone a selective sweep through *K. marxianus* populations, we could detect it across strains of this species. To test this, we carried out short-read resequencing of the genomes of five wild *K. marxianus* isolates (Table S1) and used the resulting reads alongside our reference genome of the Km31 *K. marxianus* strain as input to a pipeline for variant calling, pseudogenome construction, and sequence alignment. We then inspected these alignments at each site where the *K. marxianus* allele had been flagged as arising under selection according to our phylogenetic analysis of reference genomes. At 99% of the latter, we detected complete conservation across *K. marxianus* strains (Figure 4C and Table S5). These data establish that alleles in Km31 are representative of the population more broadly at these loci, providing further support for our inference of their derived alleles maintained under selection.

### Enrichment for amino acid variation between *Kluyveromyces* species in transporter genes

To complement our phylogenetic analyses, we next applied population-based molecular-evolution tests that used our *K. marxianus* resequencing data and a population-genomic resource for *K. lactis* (Friedrich et al. 2023). We implemented McDonald-Kreitman molecular evolution paradigms (McDonald and Kreitman 1991) to assess evidence for differences in selective pressure between the species on a given protein sequence, as follows. For each coding region we tabulated sites of non-synonymous and synonymous divergence between *K. marxianus* and *K. lactis*, and likewise for polymorphic variants within the species (Table S6).

Using these counts in a non-polarized McDonald-Kreitman test on a per-gene basis, we identified 11 genes with a significant excess of interspecies amino acid divergence across the coding region (relative to polymorphism and for synonymous variation; Figure 4D and Table S6). Thus, relative to our phylogenetic methods (Figure 4A), the McDonald-Kreitman scheme was under-powered in this system, as expected if most loci only had a few amino acid targets of adaptation between *K. marxianus* and *K. lactis* (Table S5). But among the gene hits that did emerge from the McDonald-Kreitman test, each represented an inference of positive selection from population data that complemented our phylogenetic results. Two such loci had also been top-scoring in both of our phylogenetic frameworks, the annotated glycolytic enzyme gene *FBA1* and *FAT3,* annotated in lipid transport (Figure 4C-D). Allele-swap experiments revealed a small (1.09-fold) but significant advantage accruing from the *K. lactis* allele of *FAT3* in the *K. marxianus* background in assays of lipid utilization, consonant with the better performance by wild-type *K. lactis* in the latter (Figure 4E and Figure S6); no such allele-swap effect was detectable in standard-media controls (Figure S7). This result validated our inference from sequence analyses (Figure 4C-D) of differential function of *FAT3* between the species, and attested to a modifying role for *FAT3* in a complex genetic architecture of lipid processing preferences.

For more holistic insights into *Kluyveromyces* evolution using our population-genomic data resource, we developed a group-wise version of the McDonald-Kreitman paradigm, in which we tabulated the neutrality index (NI) for a given gene as non-synonymous divergence normalized for polymorphism and for synonymous variation, and we evaluated significance of this metric across cohorts of genes of similar function via resampling (Figure 4F and Table S7). For metabolic gene groups with extreme NIs in this test, focused analyses indicated that the signal was driven by low rates of protein evolution within species, reflecting a history of purifying selection (Table S7). By contrast, for five Gene Ontology terms, the individual components of the NI metric followed the opposite pattern: non-synonymous variation between species was high in the genes of the group and polymorphism levels were on par with the genomic average (Figure 4E and Table S7). Of particular note among these gene groups enriched for low NI was a transmembrane transport gene cohort (Figure 4F), echoing our observation of membrane protein gene family expansions in *K. marxianus* (Figure 3) and the annotated transporters, including *TRK1* and *FAT3*, that had been top-scoring in our single-gene phylogenetic and McDonald-Kreitman analyses (Figure 4D and Table S5). Taken together, our molecular-evolution analyses attest to a complex history of selection on loci throughout the *K. marxianus* genome, with strong concordance across our test approaches for evidence for unique selective pressures on the coding regions of transporter genes.

## Discussion

Against the backdrop of years of work mapping the genetic basis of trait variation within fungal species (Liti and Louis 2012; Taylor et al. 2017), dissecting fixed trait divergences between species remains a key challenge for mycologists. In landmark cases, individual loci have been identified that underlie such lineage-defining phenotypes in fungi (Nagy et al. 2017; Wang et al. 2024). For the vast majority of traits varying across fungal species barriers, however, the mechanistic underpinnings remain unknown. In this work, we have established a framework for discovery of candidate genes and pathways underlying traits unique to *K. marxianus*. Our results have revealed attributes that distinguish *K. marxianus* from the rest of its genus: stress resistance in growth assays, divergent expression programs and lipid processing phenotypes, and sequence evidence for gene-content and amino-acid variation. These observations serve to complement classic trait compendia (Kurtzman et al. 2011) and the modern literature (Lane and Morrissey 2010; Lane et al. 2011; Madeira-Jr and Gombert 2018). In terms of genetics, the alleles we have found to be unique in *K. marxianus* represent candidates for the determinants that nature would have used to build the trait syndrome in this species. We expect that as these loci are validated, they will prove sufficient to reconstitute traits from *K. marxianus* in an exogenous background, as opposed to the handful of previous studies that have reported genes necessary for thermotolerance in *K. marxianus* (Falcão Moreira et al. 1998; Wu et al. 2020; Montini et al. 2022).

Most pervasive in our omic profiles was the trend of induction of membrane-gene expression, expanded membrane-gene families, and derived amino acid alleles in single-copy transporter genes, in the *K. marxianus* genome relative to the rest of its clade. The latter include gains by *K. marxianus* in sugar and potassium transporter repertoires, which were previously analyzed in comparisons with distant fungal relatives (Varela, Puricelli, Montini, et al. 2019; Papouskova et al. 2024) and which we can now date to the divergence of *K. marxianus* from its sister species. A causal phenotypic role of this remodeling of the transporter repertoire in the stress resistance by *K. marxianus* remains to be established. It is tempting to speculate that *K. marxianus* has tuned many of its transporters to tighten selectivity against extracellular toxins or to improve function in the face of heat or chemical stress. Additionally, given the evidence from our work and that of others (Dekker et al. 2021) for heightened membrane trafficking and limited exogenous lipid usage in *K. marxianus*, we propose that evolution has tuned intracellular lipid processing in this species, with molecular mechanisms to restrict exogenous uptake that involve *FAT3*. Future analyses will be necessary to test directly for a relationship between lipid processing changes and stress resistance in *K. marxianus*, potentially including specialized programs to tune membrane fluidity as seen in other yeast systems (Henderson and Block 2014; Lairón-Peris et al. 2021; Ribeiro et al. 2022; Sokolov et al. 2022). Any such unique attributes of *K. marxianus* membrane lipids could be interconnected with the genetic and expression changes we have observed at transporter genes, *e.g.* to adjust protein anchoring or to compensate for changes to membrane barrier function that would impact passive nutrient acquisition (Ferenci 2016). That said, we observed signals of evolutionary innovation by *K. marxianus* in genes of many other functions, and globally, we picture a multifaceted molecular mechanism for its resistance traits.

Why, ecologically, would *K. marxianus* have evolved stress tolerance? The ecological pressures that would have driven changes in this lineage remain poorly established. Though well-recognized as an industrial dairy yeast (Prado et al. 2015; Gethins et al. 2016; Ortiz-Merino et al. 2018; Varela, Puricelli, Ortiz-Merino, et al. 2019; Yazdi et al. 2022), *K. marxianus* was likely domesticated for this purpose fairly recently, as inferred for *K. lactis* (Friedrich et al. 2023). And though *K. marxianus* can also be isolated from rotting fruit (Williams et al. 2019; Al-Qaysi et al. 2021), it may not have specialized narrowly to such substrates: a neighboring clade that includes *Saccharomyces cerevisiae* produces more ethanol than do any of the *Kluyveromyces* species (Nespolo et al. 2020), suggesting a weaker drive in the latter toward this putative strategy to kill off microbial competitors in high-sugar environments. Among microbial-ecology findings, the strongest clue to the origin of unique phenotypes in *K. marxianus* lies in its prevalence in decomposing plant material other than fruit, including leaf litter and pine needles (Spurley et al. 2022), cow dung (Avchar et al. 2022), industrially processed cocoa beans (Baptista et al. 2021), agave (Chacón-Vargas et al. 2020), and sugar cane (Anderson et al. 1986; Cernak et al. 2018) as well as industrial and domestic compost (Suzuki et al. 2014; Avchar et al. 2022; Verma et al. 2023). Plant defense compounds in decomposing material are a compelling candidate driver for the rise of resistance traits in *K. marxianus* (Baptista and Domingues 2022) since they can persist for days in a leaf or stem after abscission (Burton 1982; Goodspeed et al. 2013; Shimada et al. 2015). Furthermore, some evidence for low pH deep in compost piles (Rainey and Covey 2016) suggests that this condition could also have contributed to *K. marxianus* evolution. In this scenario, the phenotypic syndrome of *K. marxianus* would involve not only resistance to the stresses of the decaying plant environment, but also the rapid utilization of simple sugars liberated early in plant breakdown, mediated by the new copies and new alleles of metabolic genes we have noted in the *K. marxianus* genome. As *K. marxianus* lacks the capacity to degrade lignocellulosic plant cell walls, these would be left to other fungal saprotrophs acting later in decomposition.

We thus envision an ecological model in which *K. marxianus* originally evolved as a plant decay specialist, with extant strains coming to occupy a broad range of niches, including soil and human and animal material (Rij 2013; Ortiz-Merino et al. 2018; Zhang et al. 2021). Given that we and others have seen conservation of resistance traits across *K. marxianus* strains (Lane et al. 2011; Madeira-Jr and Gombert 2018; Lappe-Oliveras et al. 2023), we infer that unique capacities arose early in *K. marxianus,* sometime after *K. marxianus* originated as a species, ∼25 million years ago (Shen et al. 2018), and persisted during its radiation into modern populations.

## Methods

### 1. Strains

Strains used in phenotyping experiments, population-genomic analyses, and mutant phenotyping are listed in Table S1.

### 2. Liquid growth phenotyping of wild-type *Kluyveromyces*

Liquid culture experiments in stress and standard-condition controls in Figure 1A used *K. marxianus* Km31 (Cernak et al. 2018); *K. lactis* yLB72 (Sorrells et al. 2015); and the strains of *K. dobzhanskii, K. wickerhamii, K. nonfermentans, K. aestuarii*, and *K. siamensis* listed in Table S1. For a given species and condition, we streaked a frozen stock onto yeast peptone dextrose (YPD) agar plates, incubated for two days at 28°C, picked a single colony, inoculated into 5 mL liquid YPD, and incubated with shaking at 28°C for 16-18 hours. We call this the first pre- culture. We diluted, into fresh liquid YPD, an inoculum of the first pre-culture at the volume required to attain an optical density (OD_600_) of 0.05 for a total culture volume of 5 mL, and we incubated this second pre-culture with shaking at 28°C for ∼7 hours until it reached OD_600_ = 0.2. For unstressed controls in Figure 1A, we diluted an inoculum of the second pre-culture into fresh liquid YPD at the volume required to attain OD_600_ = 0.2 for a total culture volume of 5 mL. 150 µL of this experimental culture was transferred to each of 12 wells of a 96-well plate, sealed with a Breathe-Easy membrane (Diversified Biotech), for incubation at 28°C and growth measurements across a 24-hour timecourse in a Tecan M200 PRO plate reader (parameter time point measurement, OD_595_). For heat stress treatment in Figure 1A, setup and growth assays were as above except that incubation of the experimental culture was at 42°C. For chemical stress treatments in Figure 1A, setup and growth assays were as above except that for the experimental culture, before transfer to the 96-well plate, ethanol (Koptec) was added to attain a concentration of 6.5% (v/v); caffeine (Sigma-Aldrich) was added to attain 25 mM; methane methylsulfonate (MMS; Thermo Scientific) was added to attain 0.03% (v/v); or propidium iodide was added to attain 1mM, and growth timecourses were 48 to 72 hours. For chemical stress treatments and growth measurements in Figures S1 and S2, for a given species and condition, streaking and the first pre-culture were as above. We diluted an inoculum of the first pre-culture into fresh liquid YPD at the volume required to attain OD_600_ = 0.2 for a total culture volume of 5 mL, with 1 M NaCl (Fisher), 3% glycerol (v/v; Fisher), 80 mM CaCl_2_ (Sigma), 250 mM rapamycin (Research Products International), 6 mM H_2_O_2_ (Fisher), or 16% (v/v) Triton X-100 (Fisher), followed by incubation and growth measurements in the 96-well format as above.

### 3. Assays of cell survival from *K. marxianus* and *K. lactis* stationary and log-phase liquid culture

For survival assays of cells in exponential phase in Figures 1B,C and S3, for *K. marxianus* Km31 and separately for *K. lactis* yLB72 we streaked a frozen stock onto yeast peptone dextrose (YPD) agar plates, incubated for two days at 28°C, picked a single colony, inoculated into 5 mL liquid YPD, and incubated with shaking at 28°C for 16-18 hours. We diluted, into fresh liquid YPD, an inoculum of this first pre-culture at the volume required to attain OD_600_ = 0.05 for a total culture volume of 5 mL, and we incubated this second pre-culture with shaking at 28°C for ∼7 hours until it reached OD_600_ = 0.2. We took this second pre-culture directly into experimental treatment as follows. For unstressed controls, serial dilutions of the culture were spotted onto agar plates and incubated for 2 days at 28°C. For heat shock, 1 mL of the culture was incubated in a heat block at 47°C for 30 minutes and then transferred to ice for 2 minutes, followed by spotting and incubation as above.

For survival assays of cells in stationary phase in Figures 1B and S3, for *K. marxianus* Km31 and separately for *K. lactis* yLB72 we streaked a frozen stock onto yeast peptone dextrose (YPD) agar plates, incubated for two days at 28°C, picked a single colony, inoculated into 5 mL liquid YPD, and incubated with shaking at 28°C for 96 hours. Heat shock and spotting (and unstressed controls) were as above.

### 4. Assays of cell survival from *K. marxianus* and *K. lactis* anaerobic and aerobic liquid cultures

As *Kluyveromyces* cannot tolerate strictly anaerobic conditions (Kiers et al. 1998; Ishtar Snoek and Yde Steensma 2006; Merico et al. 2007; Dekker et al. 2021), for Figure S4 we used a limited-oxygen experimental paradigm (Hoshida et al. 2020) as follows. For *K. marxianus* Km31 and separately for *K. lactis* yLB72, for each treatment we isolated a single colony on solid plates and carried out first and second pre-cultures in liquid YPD at 28°C under aerobic conditions as in Section 2. For unstressed controls in anaerobic conditions, we diluted an inoculum of the second pre-culture into fresh liquid YPD at the volume required to attain OD_600_ = 0.05 for a total culture volume of 5 mL. 150 µL of this experimental culture was transferred to each of 12 wells of a 96-well plate, which was placed (with lid) into an AnaeroPouch (ThermoScientific) with a AnaeroPouch - Anaero gas pack. The sealed pouch and plate were incubated for 72 hours at 28°C. Then we collected 30 µL of culture from each well and combined them all into one master culture, 100 µL of which we serially diluted into spots on agar plates, followed by incubation for 2 days at 28°C. Aerobic controls were treated analogously except that the 96-well plate was inoculated without a pouch. Viability assays of chemically stressed cells in anaerobic conditions and aerobic controls were as above except that for the experimental culture, before transfer to the 96-well plate, ethanol, caffeine, or MMS were added as in Section 2. Viability assays of heat-treated cells in anaerobic conditions and aerobic controls were as above except that the sealed pouch and plate were incubated for 24 hours at 39°C.

### 5. Transcriptional profiling

#### 5.1. Culture protocol and experimental design

For a given replicate of RNA-seq of *K. marxianus* Km31 and separately for *K. lactis* yLB72 in unstressed control conditions, we isolated a single colony on solid plates and carried out first and second pre-cultures in liquid YPD at 28°C as in Section 2. For unstressed controls, we diluted an inoculum of the second pre-culture into fresh liquid YPD at the volume required to attain OD_600_ = 0.05 for a total culture volume of 50 mL. This experimental culture was incubated at 28°C with shaking until it reached OD_600_ = 0.4-0.85. From this culture we took an aliquot whose volume contained 2 OD units of cells, pelleted its cells by centrifugation for 5 minutes at 1,500 x g, and resuspended in 1 mL of RNAlater (ThermoFisher) followed by storage at 4°C for up to 48 hours. For RNA-seq of heat-stressed cells, culture, pelleting, and storage were as above except that the experimental culture was incubated at 39°C; for RNA-seq of cells under chemical stress, culture, pelleting, and storage were as above except that the experimental culture was done in YPD with 22 mM caffeine, 0.01% MMS, or 3% ethanol. In each case, we used incubation times for the respective condition and species that allowed the culture to reach the target optical density (*i.e.* mid-exponential phase). For expression-based reannotation of *K. marxianus* Km31 and *K. lactis* yLB72 (see below), we used the above pipeline for a preliminary expression profiling experiment which we call the training batch: we cultured one biological replicate of Km31 and yLB72 at 37°C and one of Km31 and yLB72 in unstressed control conditions. For comprehensive transcriptional profiling analyses in Figure 2 and Figure S5, we used the above pipeline to generate cell pellets in two production batches. In one batch we grew three biological replicates of Km31 and three of yLB72 in 39°C heat stress, and three unstressed cultures of each species. In another batch we grew three biological replicates of each species in caffeine, three in MMS, and three in ethanol, and three unstressed controls.

#### 5.2. RNA isolation and sequencing

For RNA isolation of a given sample pellet stored in RNAlater (see Section 5.1), we centrifuged for 5 minutes at 1,500 x g, and after decanting of the supernatant, we resuspended the pellet in 200 µL of nuclease-free water followed by addition of 750 µL of BashingBead buffer from the Direct-Zol RNA MicroPrep RNA kit (Zymo Research). We then transferred our 950 µL of suspension into BashingBead tubes, vortexed for 20 minutes, and centrifuged for 5 minutes at 12,000 x g. We took ∼600 µL of supernatant from the top layer being careful not to touch the pellet, transferred to a new tube, and mixed with an equal volume of 100% ethanol. This mixture was used as input to the rest of the kit, following the manufacturer’s instructions. RNA library preparation and sequencing of ∼20M 150 bp paired-end reads per sample was carried out by Novogene, Inc.

#### 5.3. Genome re-annotation and ortholog calls for K. marxianus and K. lactis

To generate a reference genome for *K. marxianus* strain Km31, we proceeded as follows. We first used bwa (bio-bwa.sourceforge.net) to map the RNA-seq reads from the *K. marxianus* cultures of the training batch of our transcriptional profiling (see Section 5.1) to a published genome of the DMB1 strain of *K. marxianus* (Suzuki et al. 2014). We then used these mappings, requiring mapping qualities of >20 and mapped lengths >20, to call single-nucleotide polymorphisms with BCFtools (Li 2011; Danecek et al. 2021) mpileup with the -I flag to ignore indels, and filtering for exclusion of variants with: a depth of less than 10 reads, or more than 10% of reads with a mapping quality of 0 supporting the variant. At the called polymorphisms, we computationally introduced Km31 alleles into the DMB1 genome sequence with the consensus tool in BCFTools to generate a first-pass pseudogenome for Km31. Next, we used this pseudogenome and the training batch RNA-seq reads as input into transcript assembly with Trinity (Grabherr et al. 2011) and annotation with PASA (Haas et al. 2003), and we took the output as our final annotated genome for *K. marxianus* Km31 (Supplementary Data 1). We generated a reference genome for *K. lactis* strain yLB72 as above, except that we used RNA-seq from the *K. lactis* training batch and the published genome of *K. lactis* strain NRRL Y-1140 (Dujon et al. 2004) (Supplementary Data 1). To call orthologous gene pairs between *K. marxianus* Km31 and *K. lactis* yLB72, we used our final annotations from the two species as input into OrthoFinder (Emms and Kelly 2019). We retained all one-to-one orthology calls, *i.e.* cases in which a single annotated gene from *K. marxianus* was called orthologous to a single gene from *K. lactis*. We also sought to valorize cases in which OrthoFinder, with the goal of finding orthologs in species B of a gene *G1,A* in species A, did not distinguish between *n* candidate orthologous genes *G1…n,B* from species B. We evaluated each candidate *Gi,B* for synteny: if the next gene up- or downstream of *Gi,B* in the species B genome was called a one-to-one ortholog of the up- or downstream genes of *G1,A* in the species A genome, we retained this case as a bona fide ortholog pair. Our final set comprised 3578 ortholog pairs between *K. marxianus* Km31 and *K. lactis* yLB72. *Saccharomyces* orthologs were taken from (Sorrells et al. 2018).

#### 5.4. Production-batch transcriptional profiling data analysis

For comprehensive transcriptional profiling analyses in Figure 2 and Figure S5, we used our RNA-seq data from cultures of *K. marxianus* Km31 and *K. lactis* yLB72 from production batches (see Section 5.1) as follows. For the RNA-seq reads from a given replicate of a given species (*K. marxianus* or *K. lactis*) and condition (39°C, ethanol, caffeine, MMS, or unstressed control), we mapped to the respective pseudogenome (see Section 5.3) using the TopHat (Kim et al. 2013) aligner with default parameters, and we used these mappings as input into HTseq (Anders et al. 2015) to generate raw read counts per gene. For a given species and stress, we used the counts from all replicates from the stress, and the unstressed control from the respective batch, as input into edgeR (Robinson et al. 2010) to generate normalized counts per gene and replicate and a normalized log_2_ fold-change between stress and control conditions as an average across replicates. Principal component analysis (PCA) of the RNA-seq data was done using the PCA module of scikit-learn (Pedregosa et al. 2011). The edgeR normalized counts for each gene in each treatment were scaled to unit variance with scikit-learn before running the PCA. Principal component weights are listed in Supplementary Table 2.

To identify Gene Ontology (GO) terms associated with principal components, we assigned GO annotations to each gene using the *K. lactis* annotation from FungiDB (https://fungidb.org/fungidb/app/record/dataset/NCBITAXON_284590). For each GO term with at least three annotated genes in our dataset, we tested whether genes within that term had significantly extreme PC1 or PC2 weights compared to the null expectation. We used a bootstrap resampling approach to generate the null distribution: for each GO term containing n genes, we randomly sampled n genes (with replacement) from the complete set of genes and calculated the mean PC weight of this random sample. This process was repeated 10,000 times to construct an empirical null distribution representing the expected mean PC weight for a random gene set of that size. We calculated two-tailed *p*-values by comparing the observed mean PC weight to this null distribution. To correct for multiple testing across all GO terms, we applied the Benjamini-Hochberg false discovery rate (FDR) correction using the multipletests function from statsmodels (Seabold and Perktold 2010), with a significance threshold of α = 0.05.

To identify gene groups in which a given stress treatment evoked significant gene induction or repression relative to the unstressed control, in *K. marxianus* Km31 and separately in *K. lactis* yLB72 (Table S3), we carried out resampling-based enrichment tests as follows. We used assignments of each gene to Gene Ontology terms for *K. lactis* from FungiDB (https://fungidb.org/fungidb/app/record/dataset/NCBITAXON_284590), and we propagated these assignments to orthologous genes in *K. marxianus* from Section 5.3. For a given species and stress, for each Gene Ontology term in turn, for the *n* genes from the term with expression observations, if *n* > 5 we collated the genes’ log_2_ fold-changes in expression between stress and the untreated control, each of which was a mean across replicates (from edgeR; see above). We took the mean of these fold-changes across the genes of the term, *mtrue*. We then picked *n* random genes from the genome and tabulated their mean expression fold-change for the respective species and stress, *mresample*. Repeating the latter resampling 1000 times, we calculated an empirical *p*-value reporting the enrichment of the magnitude of expression fold-change as the percentage of resamples for which (|*mresample*| ≥ (|*mtrue*|). We used the *p*-values for all terms tested for the respective species and stress as input into multiple testing correction with the Benjamini-Hochberg method.

### 6. Molecular-evolution analyses

#### 6.1. Gene duplication analysis

As input to the gene family copy number analysis for Figure 3 and Table S4, genomes and annotations were as follows. For *K. marxianus* and *K. lactis* we used the reference genomes and annotations of Km31 and yLB72, respectively, from Section 5.3. For *K. wickerhamii* we used the reference genome and annotation of UCD54-210 from (Sorrells et al. 2018). For the *K. nonfermentans* strain NRRL Y-27343 (GenBank acc. no. GCA_030569915.1), the *K. aestuarii* strain NRRL YB-4510 (GenBank acc. no. GCA_003707555.2), the *K. siamensis* strain CBS 10860 (GenBank acc. no. GCA_030579315.1), and the *K. dobzhanskii* strain NRRL Y-1974 (GenBank acc. no. GCA_003705805.3) we used YGAP (Proux-Wéra et al. 2012) to generate *de novo* annotations from previously generated reference genomes (Supplementary Data 1). As outgroup species we used the genomes and annotations for *Ashbya aceri* (GenBank acc. no. GCA_000412225.2), *Candida albicans* (GenBank acc. no. GCA_000182965.3), *Eremothecium cymbalariae* (GenBank acc. no. GCA_000235365.1), *Eremothecium gossypii* (GenBank acc. no. GCA_000091025.4), *Eremothecium sinecaudum* (GenBank acc. no. GCA_001548555.1), *Lachancea dasiensis* (GenBank acc. no. GCA_900074725.1), *Lachancea fantastica* (GenBank acc. no. GCA_900074735.1), *Lachancea fermentatii* (GenBank acc. no. GCA_900074765.1), *Lachancea lanzarotensis* (GenBank acc. no. GCA_000938715.1), *Lachancea meyersii* (GenBank acc. no. GCA_900074715.1), *Lachancea mirantina* (GenBank acc. no. GCA_900074745.1), *Lachancea nothofagi* (GenBank acc. no. GCA_900074755.1), *Lachancea quebecensis* (GenBank acc. no. GCA_002900925.1), *Lachancea thermotolerans* (GenBank acc. no. GCA_000142805.1), and *Saccharomyces cerevisiae* (GenBank acc. no. GCA_000146045.2). We used proteomes for each species as input into OrthoFinder (Emms and Kelly 2019) to find groups of orthologs. We then used the time-weighted tree and gene family counts from OthoFinder as input into CAFE 5 (Mendes et al. 2021) to analyze gene family contractions and expansions. Gene families that did not exist at the root were included. We adjusted *p*-values with the Benjamini-Hochberg test to correct for multiple testing. We used CafePlotter (https://github.com/moshi4/CafePlotter) to visualize significant gene family expansions and contractions on the phylogenetic tree. We then assigned functions for each gene family by running all proteins from all species for each gene family through InterProScan (Jones et al. 2014). We took the most common InterPro term in the output of the scan as the inferred function of the gene family in Figure 3.

#### 6.2. Phylogenetic analyses

As input to phylogenetic analyses of one strain for each *Kluyveromyces* species for Figure 4 and Table S5, we used genomes and annotations for the *Kluyveromyces* genus excluding *K. siamensis* as in Section 7.1. We called orthologs across these six species with OrthoFinder (Emms and Kelly 2019), and for a given orthogroup we generated protein alignments using MUSCLE (Edgar 2004). We then used this alignment and its nucleotide sequences as input into PAL2NAL (Suyama et al. 2006) using the -nogap flag to remove gaps to generate a codon alignment. For each orthogroup in turn, we ran the PAML (Yang 2007) codeml package run using the branch-site model (Zhang et al. 2005) to evaluate the fit to the sequence data of a model in which regions of the coding sequence are under positive selection for amino acid variation along a focal (foreground) branch and not on any other branch from the tree (the background branches). We used as input to PAML the codon alignment for the orthogroup and the topology of the phylogenetic tree relating the species (as in Figure 1A), omitting *K. siamensis*. The calculation proceeded in two steps. First, we fit a null model in which all branches of the tree had the same protein evolutionary rate (PAML codeml run with options verbose = 1, seqtype = 1, clock = 0, model = 2, NSsites = 2, fix_omega = 1, omega = 1), yielding a protein evolutionary rate and log-likelihood of the data under the model. Next, we re-fit using an alternative model in which the branch leading to *K. marxianus* was the foreground (PAML codeml run with options verbose = 1, seqtype = 1, clock = 0, model = 2, NSsites = 2, fix_omega = 0). Outputs were the codon sites assigned to the class with an excess of amino acid changes in *K. marxianus* relative to silent changes, and their posterior probabilities under Bayes Empirical Bayes inference (Yang et al. 2005); the codon sites assigned to classes with relaxed and purifying selection, respectively, along the background lineages, and their posterior probabilities; and the log-likelihood of the data under the model. The *p*-value assessing the improvement in fit to the data of the alternative model relative to the null was calculated from a likelihood ratio test (a χ^2^ test on twice the difference of likelihoods with one degree of freedom).

*p*-values were corrected for multiple testing by the Benjamini/Hochberg method using the multipletests function from statsmodels (Seabold and Perktold 2010). In Figure 5 A we tabulated all genes with adjusted *p* < 0.05 from this test that had more than 1% of codon sites assigned to the class with an excess of amino acid changes in *K. marxianus* (proportion in class 2a > 0.01 in codeml). Separately, we repeated the above analyses using the branch leading to *K. lactis* as the foreground. Conservation of sites called significant by PAML using *K. marxianus* as the foreground were calculated using Biopython (Cock et al. 2009) to count the conservation of each site among our sequenced *K. marxianus* strains (Table S1 and see Section 7.3). *Saccharomyces* orthologs were taken from (Sorrells et al. 2018).

To complement our PAML analysis, we also ran HyPhy aBSREL (Pond et al. 2005) on the PAL2NAL alignments across *Kluyveromyces* orthologs as described above. aBSREL uses a branch-site model similar to PAML to detect positive selection on individual branches. We ran aBSREL indicating either *K. marxianus* or *K. lactis* as the foreground. We pulled the *p*-value as calculated by aBSREL for each orthogroup and corrected for multiple testing by the Benjamini/Hochberg method using the multipletests function from statsmodels (Seabold and Perktold 2010). *Saccharomyces* orthologs were taken from (Sorrells et al. 2018). We compared HyPhy aBSREL hits against PAML branch-site hits to confidently list genes acted on positive selection in *K. marxianus*.

#### 6.3. Resequencing and analysis of K. marxianus and K. lactis population genomes

For resequencing, we cultured, in 10 mL of liquid YPD, each of five *K. marxianus* strains of non-domesticated provenance beside Km31, and two non-domesticated *K. lactis* strains beside yLB72 (Table S1), and we isolated DNA from each with the Quick-DNA^TM^ Fungal/Bacterial Miniprep Kit (Zymo Research). Genomic DNA sequencing libraries were made and sequenced on an Illumina NovaSeq 6000 by the UC Berkeley Vincent J. Coates Genomic Sequencing Laboratory. We used Minimap2 (Li 2018) to map the *K. marxianus* reads to our *K. marxianus* Km31 reference genome (see Section 5.3). Separately, we mapped the reads from each of 41 *K. lactis* strains previously sequenced (Friedrich et al. 2023), and those from our two sequenced *K. lactis* strains, to our *K. lactis* yLB72 reference genome. For each strain we sorted mappings using the SAMtools fixmate and markdup functions, and we called single-nucleotide polymorphisms and generated a pseudogenome as in Section 5.3, except the variant filtering included only a quality score (QUAL) threshold of above 20. Called variants are provided in Supplementary Data 2. For each pair of orthologous genes between *K. marxianus* and *K. lactis* from Section 5.3, we aligned the nucleotide sequences of the coding regions from all strains of the two species using MUSCLE (Edgar 2004). We then used this alignment and its associated nucleotide sequences as input into PAL2NAL (Suyama et al. 2006) using the -nogap flag to remove gaps to generate a codon alignment.

For Figure 4D,E and Table S6,7 we carried out per-gene McDonald-Kreitman analyses (McDonald and Kreitman 1991) on the PAL2NAL alignments using the mktest function from the CodonAlignment class from Biopython. *p*-values were corrected for multiple testing by the Benjamini/Hochberg method using the multipletests function from statsmodels (Seabold and Perktold 2010). For a given gene, this approach tabulates the rates of non-synonymous polymorphism within species (P_n_), synonymous polymorphism (P_s_), non-synonymous divergence between species (D_n_), and synonymous divergence (D_s_), and the neutrality index NI = (P_n_/P_s_)/(D_n_/D_s_). Because low counts of amino acid polymorphism can drive significance in the McDonald-Kreitman framework, we developed a filtering scheme in which we tabulated the average *a* and standard deviation σ of P_n_/P_s_ across all genes for which we had sequence data. We only retained for Figure 5D and NI genomic analyses those genes with P_n_/P_s_ > *a* – 1.5σ.

For genomic analyses of P_n_/P_s_ and D_n_/D_s_ in the McDonald-Kreitman framework in Table S7, we used Gene Ontology term assignments and resampling (using 10,000 resamples) as in Section 5.4, except that we only included Gene Ontology terms containing 10 genes or more after filtering, and for P_n_/P_s_ we tested for enrichment of low values of the metric across a given term. Genomic analyses on NI proceeded as above, except on the filtered dataset of genes with P_n_/P_s_ > *a* – 1.5σ as explained above and testing for enrichment of low values of the metric across the term. *p*-values were corrected for multiple testing by the Benjamini/Hochberg method using the multipletests function from statsmodels (Seabold and Perktold 2010).

## 7. *FAT3* allelic swap mutant generation and cerulenin with oleic acid growth phenotyping

To search for *K. marxianus* phenotypes that *FAT3* could be implicated in, we assayed the genus in media with cerulenin and oleate according to the Tecan 96-well plate experimental design described in Section 2, except media contained 500 µM oleate (Sigma) and 0.5% Tween 40 (Sigma) with or without 25 µM cerulenin (from 20 mg/mL stock in ethanol; Cayman Chemical). We gathered final OD readings as an efficiency read out and tested for significance with a Mann-Whitney U test from SciPy.

We made Δ*DNL4* mutants of *K. marxianus* Km31 and *K.* lactis yLB72 as described in (Rajkumar and Morrissey 2022). We then made CRISPR/Cas9 plasmids to target each species’ *FAT3* allele in the Δ*DNL4* mutants. We chose gRNAs for *K. marxianus* based on the dataset generated from sgRNAcas9 (Xie et al. 2014) in (ref above), and chose gRNAs for *K. lactis* using CCTop (Stemmer et al. 2015). Alongside the CRISPR plasmids we made repair fragments by PCR, with 1kb regions of the species’ native gDNA flanking the opposite species’ *FAT3* allele to allow for homologous recombination. We transformed the stitched amplicon and CRISPR plasmid into *K. marxianus* and *K. lactis* ΔDNL4 mutants by electroporation (Hsu et al. 2021) and selected transformants by plating onto agar plates with 50 μg/mL of Hygromycin B (Invitrogen by Thermo Fisher Scientific). Nanopore sequencing of the ∼4kb region including the swapped allele, the flanking homology arms, and the genomic region around it confirmed positive swaps.

We assayed the growth in cerulenin and oleate of *K. marxianus* Km31, *K. lactis* yLB72, and the *FAT3* allelic swaps in either direction to assess the efficiency of *FAT3*. We streaked out *K. lactis* (Δ*DNL4*), *K. marxianus* (Δ*DNL4*), two replicate clones of *K. lactis* (Δ*DNL4*; Δ*FAT3*::*K. marxianus FAT3*), and two replicate clones of *K. marxianus* (Δ*DNL4*; Δ*FAT3*::*K. lactis FAT3*) from -80°C frozen stocks onto YPD plates and incubated them at 28°C for 2 days to allow for colony formation. We inoculated one colony per strain into 3 ml of YPD and incubated at 28°C overnight. We then back-diluted the cultures into 10 ml of YPD to achieve 0.05 OD_600_/ml. We let the cultures grow at 28°C with shaking for 6-7 hours to reach the early log-phase (0.2-0.4 OD_600_). We back-diluted the early log-phase cultures into 3 ml of oleate media (YPD with 0.5% Tween-40, 0.7% KH_2_PO_4_, 25 μM cerulenin, and 500 μM oleate) or control media (YPD with 0.5% Tween-40 and 0.7% KH_2_PO_4_). We incubated the cultures at 28°C with shaking for 24 hours and recorded the OD600 as the final readout for cell density. We ran three technical replicates per assay with at least 4 independent assays, from streaking to measurements of OD_600_.

## Acknowledgements

The authors thank Adam Arkin for his generosity with computational resources; Jamie Cate and Alexander Johnson for strains; and Jacob Steenwyk, Jeremy Roop, and members of the Brem lab for helpful discussions. This work was supported by NIH 2R01GM120430 to R.B.B. and an NSF Graduate Research Fellowship to A.D.

## Data Availability

Raw RNA-seq reads are available on GEO under series record GSE302710. Genome sequencing data is available on SRA with accession PRJNA1291269.

