## Supplemental Figures and Table Captions for "Signatures of innovation and selection in the extremotolerant yeast *Kluyveromyces marxianus*"

**Figure S1**

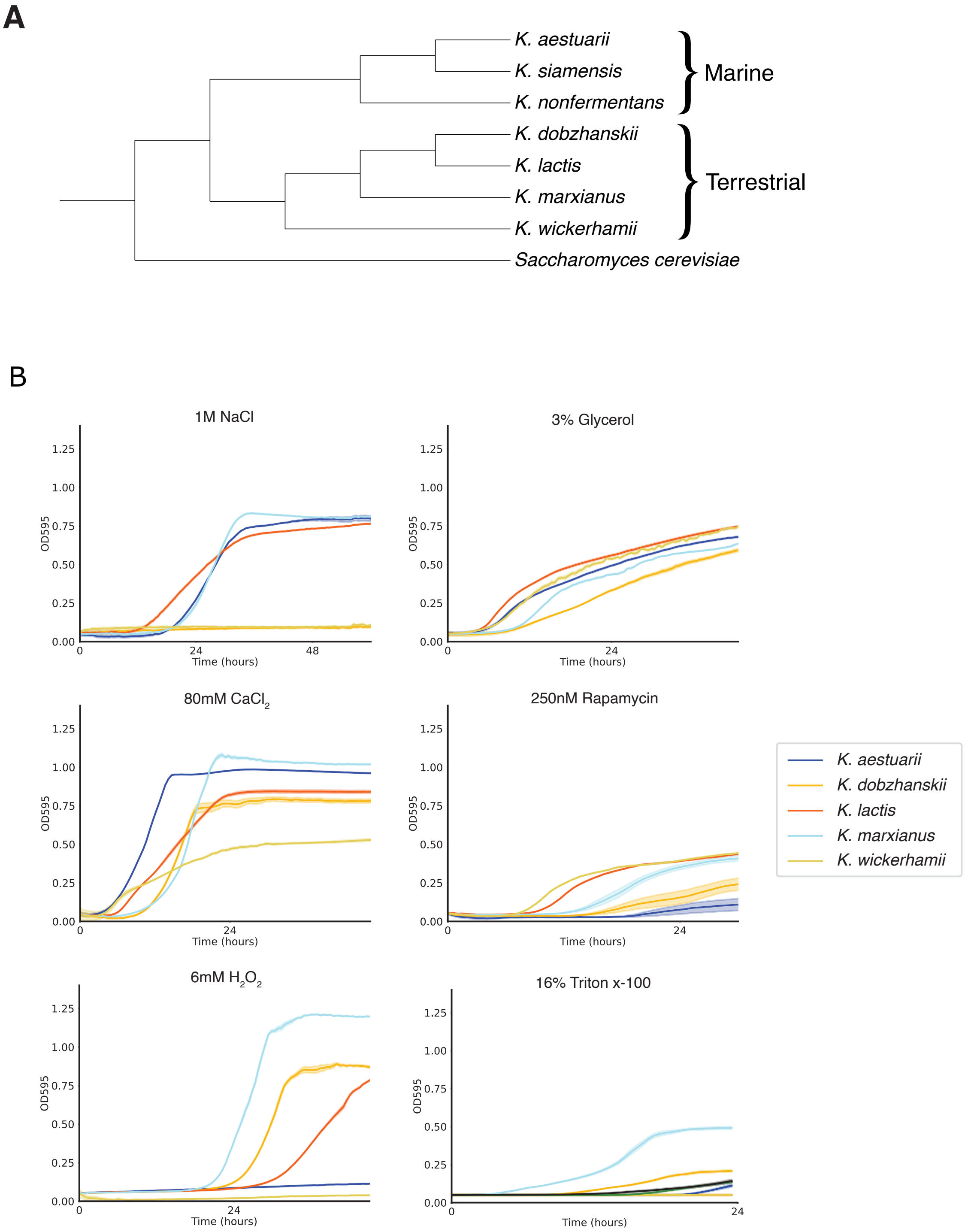


**Figure S1. Stresses in which *K. marxianus* does not exhibit unique stress tolerance relative to the rest of the genus**. **A**, Phylogenetic tree of the *Kluyveromyces* genus (Am-In et al. 2008; Varela, Puricelli, Ortiz-Merino, et al. 2019). Marine and terrestrial species are delineated according to species descriptions (Rij 2013). Branch lengths are not to scale. **B**, Data and symbols are as in Figure 1B of the main text except that each panel reports growth measurements in the indicated stress.

**Figure S2**

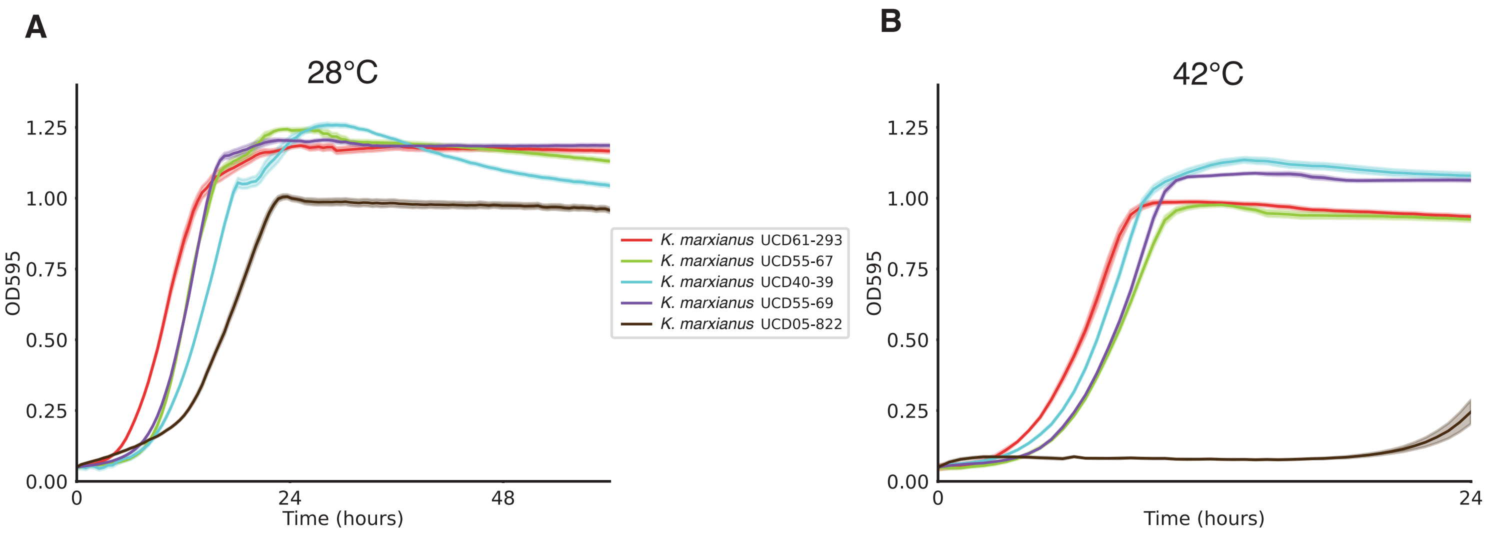


**Figure S2. Heat tolerance of *K. marxianus* strains.** Data and symbols are as in Figure 1A of the main text except that in a given panel, each trace reports growth measurements of the indicated wild K. marxianus strain.

Figure S3

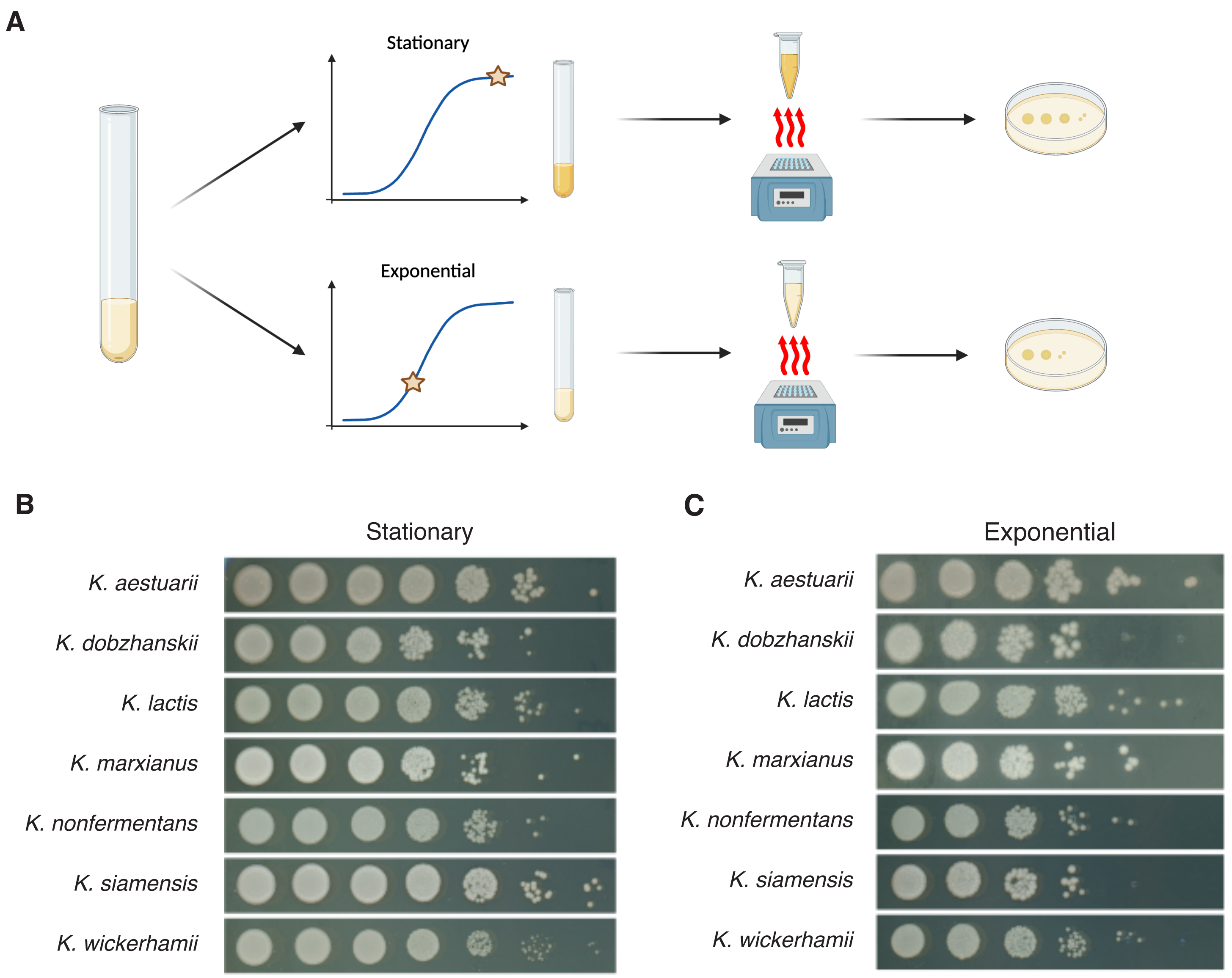


**Figure S3. No detectable difference between *Kluyveromyces* species with respect to cell viability from unstressed liquid cultures. A**, Experimental workflow created with BioRender.com. From left, inoculum from a starting liquid culture is introduced into fresh liquid medium for regrowth at 28°C and then sampled at nutrient exhaustion (Stationary) or in log phase (Exponential). The liquid sample is subjected to a heat shock of 47°C for 30 minutes and then spotted as serial dilutions onto solid medium. Viable cells after heat shock grow into colonies after incubation at 28°C. **B**, Data and symbols are as in Figure 1B of the main text except that liquid cultures were not subjected to heat shock before plating.

#### Figure S4

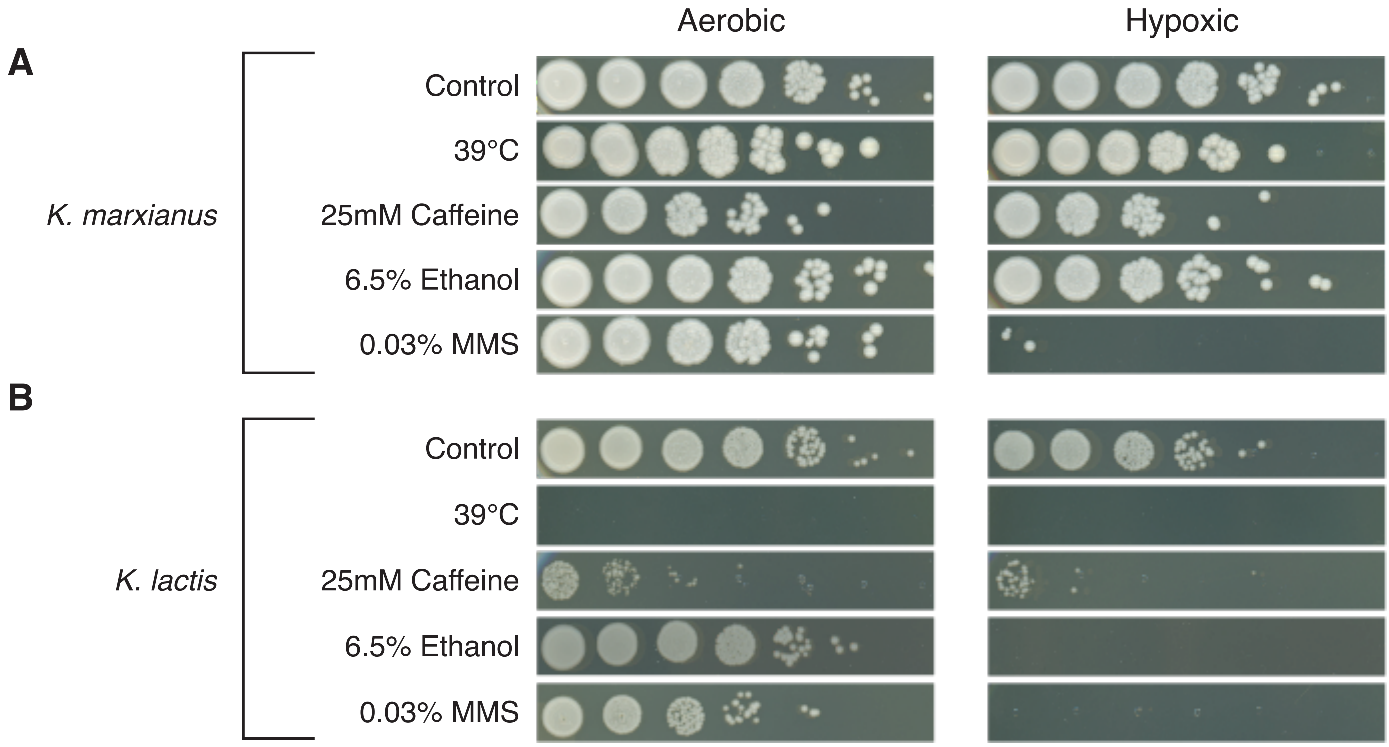


**Figure S4. Culture-based block of respiration compromises stress tolerance in *K. marxianus* and *K. lactis*. A**, **B**, Each panel reports cell viability assays after culture in liquid medium with stress, or an unstressed control, of *K. marxianus* or *K. lactis* in normoxia or hypoxia. In a given panel, each row shows serial dilutions of the liquid culture spotted onto solid medium and incubated without stress at 28°C.

#### Figure S5

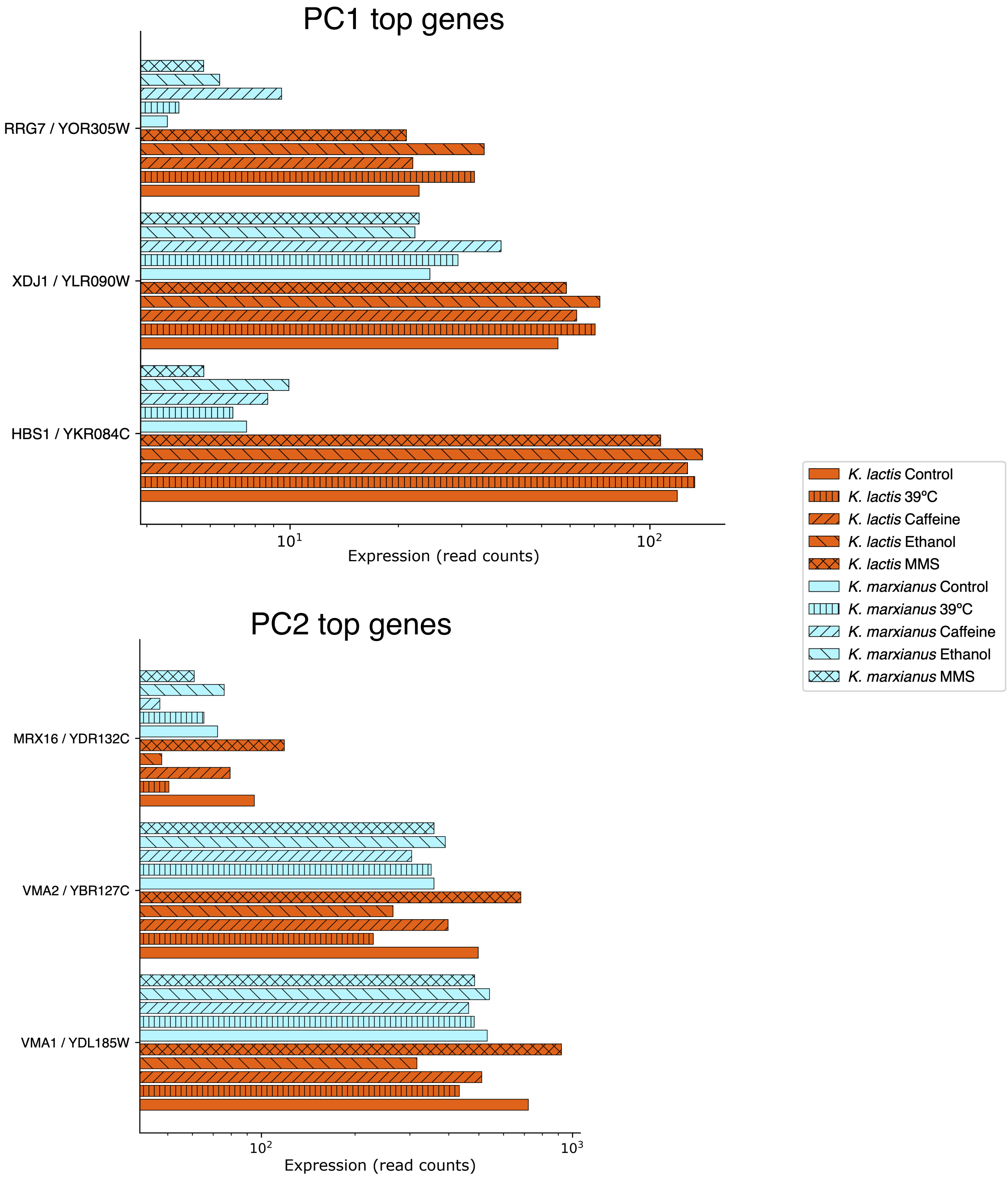


**Figure S5. Example genes contributing to principal components explaining expression divergence between *K. lactis* and *K. marxianus*. A**, Shown are results from the principal component analysis of *K. lactis* and *K. marxianus* transcriptomes across treatments in Figure 2A of the main text. Each panel reports expression of the three genes contributing most to the indicated principal component (PC1 explaining primarily differences between species in all conditions, including control, and PC2 explaining differences between treatments, primarily in *K. lactis*). In a given panel, each bar reports expression, as a mean across replicates, of the indicated gene in the indicated species and treatment (orange, *K. lactis*; blue, *K. marxianus*). The full set of gene contributions to principal components is listed in Supplementary Table 2.

### Figure S6

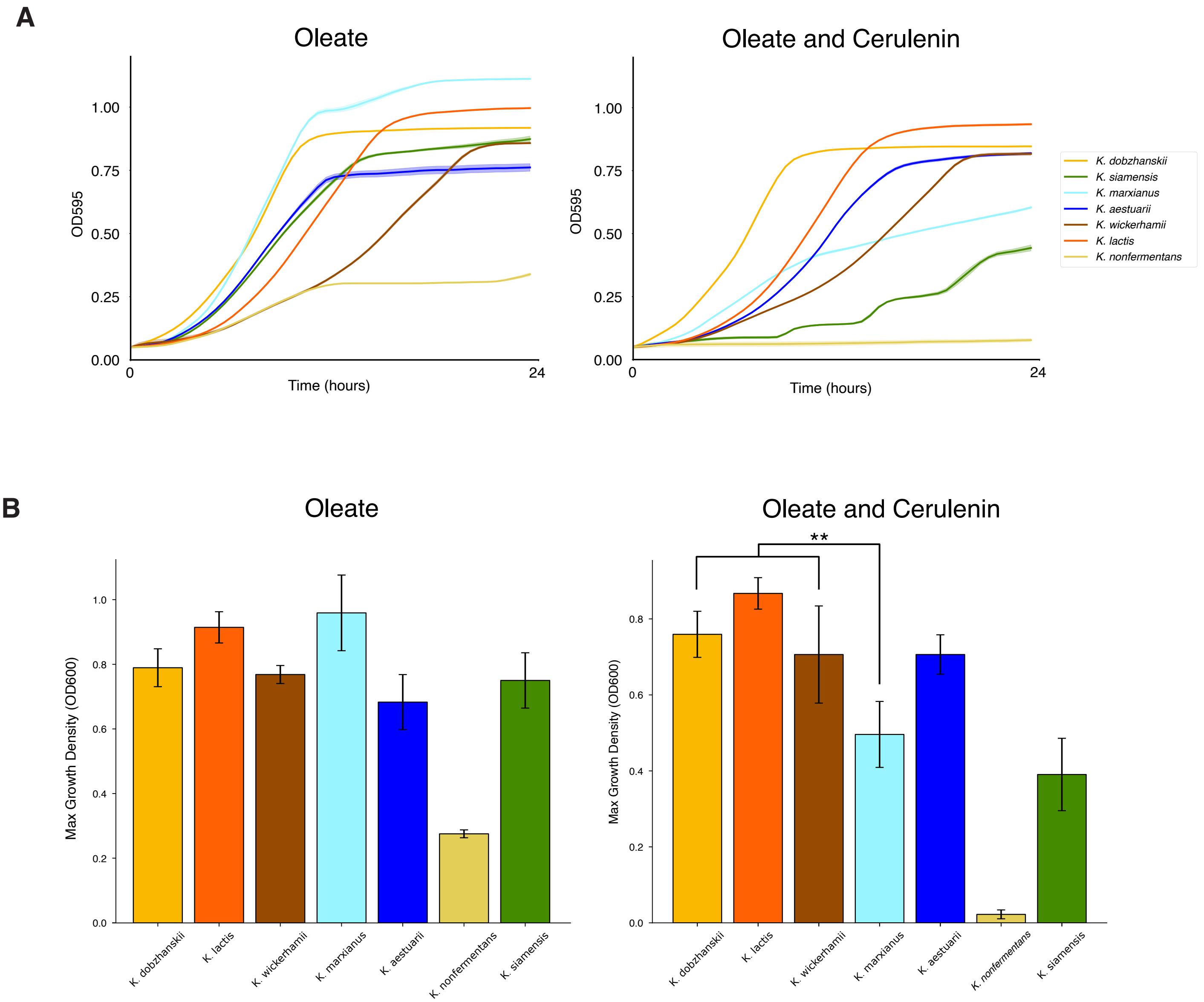


**Figure S6. *K. marxianus* shows decreased growth in cerulenin and oleate. A**, Growth curves of *Kluyveromyces* species grown with oleate with and without the fatty acid synthesis inhibitor cerulenin. Each trace reports a growth timecourse for the indicated species, with the solid line displaying the mean across technical replicates and the faint outline showing the standard error. **B**, Each bar shows the final OD reached for each *Kluyveromyces* species with oleate utilization ability after three independent runs under the same conditions as A. The starred comparison between *K. marxianus* and the other species indicate a statistically significant defect ($p = 0.0091$).

**Figure S7**

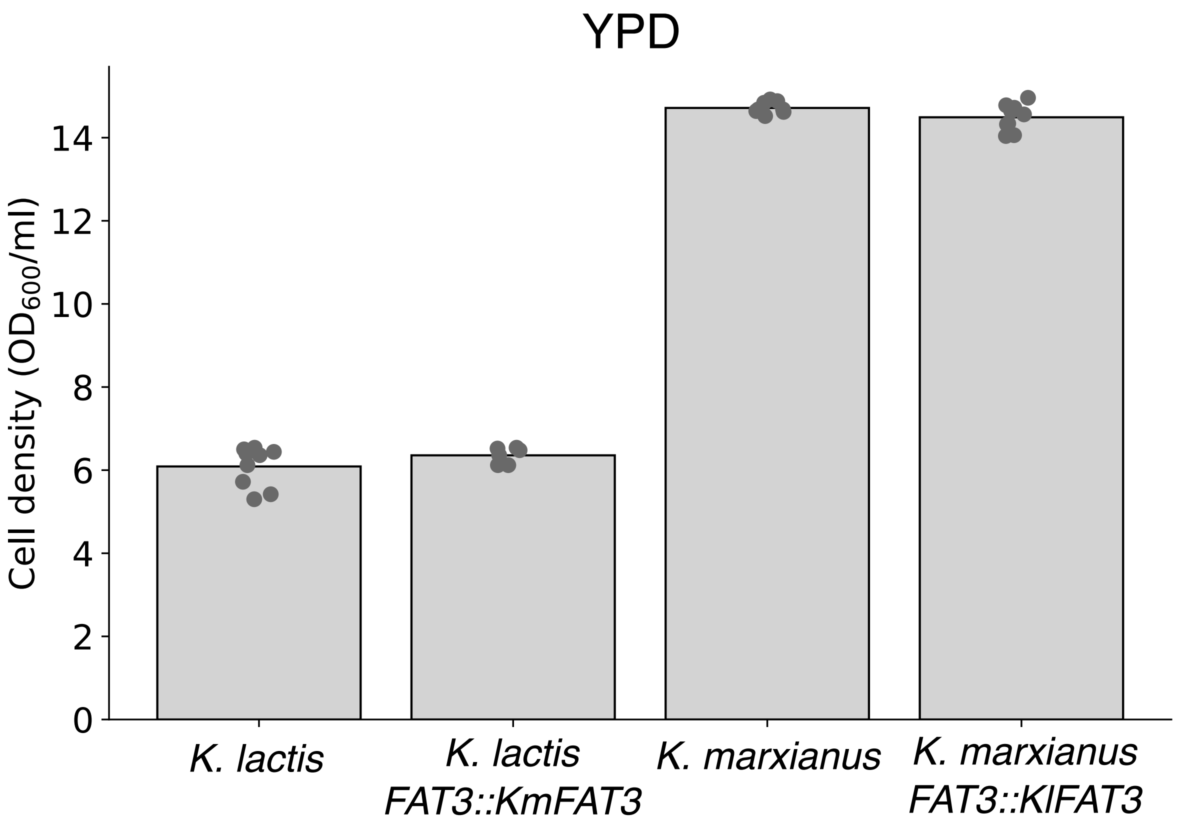


**Figure S7. Control growth of *FAT3* allelic swaps do not show same growth differences as in oleate and cerulenin.** Data are as in Figure 4E except growth is under control YPD conditions.

#### Supplementary table captions

**Table S1. Strains used in this work.** List of strains used for phenotypic assays and population analyses. All species shown are haploid.

**Table S2. Normalized RNA-seq read counts, major principal component weights, and stress effects.** Shown are details from the RNA-seq and principal component analysis of Figure 2 of the main text. Each row in Tab 1 reports RNA-seq results and analysis for a pair of orthologous *K. lactis* and *K. marxianus* genes. Columns B and C report respective weighting of the gene in the first and second components of a principal component analysis. In columns D-S, each column reports normalized RNA-seq read counts as an average across replicates from cultures of *K. lactis* or *K. marxianus* in unstressed control conditions (LC and MC, respectively) or in stress treatments (LT or MT, respectively), where stresses are 39°C, ethanol (ET), MMS, or caffeine (CAF). The next eight columns report the effect of stress on expression; each reports the average across replicates of $\log_2$(treatment/control) for each species in the indicated treatment. The last two columns list the S. cerevisiae ortholog and annotated function. Tab 2 reports results of Gene Ontology enrichment tests on PC1 weights. For each GO term, the number of genes analyzed, observed mean of PC1 weights for genes in that term, expected mean of PC1 weights, p-value, p-value adjusted for multiple testing, and effect size of the PC1 weight mean is reported. In column D, for genes that contribute to the principal component with a negative weight (blue), high expression in a given sample drives a negative PC1 value, as seen for *K. marxianus* transcriptomes (Figure 2A of the main text); likewise, for genes that contribute with a positive weight (orange), high expression in a given sample drives a positive PC1 value, as seen for *K. lactis* transcriptomes. Tab 3 reports Gene Ontology enrichment test results as in Tab 2, except that PC2 weights were analyzed. Tab 4 lists the GO term assigned to each pair of orthologous *K. lactis* and *K. marxianus* genes, along with the *S. cerevisiae* ortholog and function.

**Table S3. Gene Ontology terms with stress-responsive expression.** Each tab reports the results of tests of Gene Ontology terms for enrichment of expression change between the indicated stress and unstressed controls in the indicated species. In a given tab, each row reports results for the indicated Gene Ontology term: the average, across genes and replicates, of $\log_2$(treatment/control), and the multiple testing-corrected significance of the enrichment for high magnitude of this value relative to a resampling null.

**Table S4. Gene family analysis for expansion/contraction analysis.** Tab 1 reports the results from the CAFE analysis for gene family copy number change. Column B denotes the *p*-value from CAFE for a change in copy number in *K. marxianus* of the orthogroup indicated in Column A. Column C shows the *p*-value after adjustment for multiple testing. Tab 2 shows the genes called in each orthogroup for all species analyzed in CAFE.

**Table S5. Phylogenetic tests for evidence of positive selection.** In a given row in Tab 1, columns C-I report the results of PAML branch-site tests package for positive selection in *K. marxianus* on the indicated orthogroup (see Tab 4 for full orthogroup membership). Columns C-D report the significance of the improvement in fit to an alternative model stipulating positive selection relative to a null without positive selection, as a nominal *p*-value and after adjustment for multiple testing. Columns E and G report the number and proportion, respectively, of sites inferred to be under positive selection in the alternative model. Column F reports the average conservation of the latter across *K. marxianus* population isolates from Supplementary Data 2.

Columns H and I report inferred protein evolutionary rates for the branch leading to *K. marxianus* (foreground) and for all other branches of the tree (background), respectively. Columns J-O report the analogous results from tests for positive selection in *K. lactis*. Columns P-Q list the gene name and function for the *S. cerevisiae* ortholog. Green highlighted cells indicate cases of $p < 0.05$. For each row in Tab 2, columns B-C report the *p*-value of HyPhy aBSREL tests for positive selection in *K. marxianus* and *K. lactis* on the indicated orthogroup (see Tab 4 for full orthogroup membership). Columns D-E report the *p*-value after adjusting for multiple testing, with green columns indicating cases of $p < 0.05$. Tab 3 shows the genes listed as significant in both PAML and HyPhy tests, with columns C-G reporting PAML data as shown in Tab 1. Columns H-I list the gene name and function for the *S. cerevisiae* ortholog. Tab 4 shows the genes in each orthogroup analyzed in PAML and HyPhy, with each row reporting the genes of the indicated species representing the members of one inferred orthogroup.

**Table S6. McDonald-Kreitman tests for evidence of positive selection.** Each row reports results from a McDonald-Kreitman test on the indicated pair of orthologous genes between *K. lactis* and *K. marxianus*. $D_s$, number of divergent synonymous changes between species; $D_n$, number of divergent non-synonymous changes between species; $P_s$, number of polymorphic synonymous changes within species; $P_n$, number of polymorphic non-synonymous changes within species; NI, neutrality index (see Methods); *p* and *p*-value, nominal and multiple testing-corrected McDonald-Kreitman significances, respectively. Columns L-M list the gene name and function for the *S. cerevisiae* ortholog. Green highlighted cells indicate cases of $p < 0.05$.

**Table S7. Gene Ontology terms with evidence for positive or purifying selection.** Each tab reports the results of tests of Gene Ontology terms for enrichment of extreme values of one sequence-based metric of selection. These tests used the gene-based variation data from Table

S5 and were, respectively, for low values of the neutrality index, NI, indicative of positive selection; for low values of $P_s/P_n$, indicative of purifying selection; and for high values of the protein evolutionary rate $D_s/D_n$, indicative of positive selection. In a given tab, each row reports results for the indicated Gene Ontology term: the average across genes of the indicated metric, and the nominal and the multiple testing-corrected significance of the enrichment for extreme magnitude of this value relative to a resampling null (p-value and p_adj, respectively).
